# The structural basis of the Talin-KANK1 interaction that coordinates the actin and microtubule cytoskeletons at focal adhesions

**DOI:** 10.1101/2023.02.23.529676

**Authors:** X Li, B.T Goult, C Ballestrem, T Zacharchenko

## Abstract

Adhesion between cells and the extracellular matrix (ECM) is mediated by heterodimeric (αβ) integrin receptors that are intracellularly linked to the contractile actomyosin machinery. One of the proteins that control this link is talin, which organises cytosolic signalling proteins into discrete complexes on β-integrin tails referred to as focal adhesions (FAs). The adapter protein KANK1 binds to talin in the region of FAs known as the adhesion belt. Here, we developed a novel crystallographic method to resolve the talin-KANK1 complex. This structure revealed that the talin binding KN motif of KANK1 has a novel fold, where a β-turn stabilises the α-helical region, explaining its specific interaction with talin R7 and high affinity. Single point mutants in KANK1 identified from the structure abolished the interaction and enabled us to examine KANK1 enrichment in the adhesion belt. Strikingly, in cells expressing a constitutively active form of vinculin that keeps the FA structure intact even in the presence of myosin inhibitors, KANK1 localises throughout the entire FA structure even when actomyosin tension is released. We propose a model whereby actomyosin forces on talin eliminate KANK1 from talin binding in the centre of FAs while retaining it at the adhesion periphery.

## Introduction

The adhesion of cells to the extracellular matrix (ECM) controls cell migration, proliferation and differentiation ^[1–3]^. The cytoplasmic adapter protein talin controls the ability of cells to adhere to the ECM. Intracellular binding of talin to integrin adhesion receptors activates them and initiates the formation of adhesion complexes, that upon linkage to the force-inducing actomyosin cytoskeleton mature into larger cell-ECM contacts known as focal adhesion (FAs)^[2, 4]^. KANK proteins (isoforms 1-4) are known to bind to talin, but unlike talin, which is ubiquitous throughout the FA they localise to a belt region in the periphery of FAs ^[5, 6]^. Here they recruit the cortical microtubule stabilising complex (CMSC) formed of α and β liprins, LL5β and KIF21A, which organises microtubule plus ends at the cell cortex ^[6, 7]^. More detailed mechanistic insight into KANK recruitment and localisation requires structural insight, however, the precise structural determinants of this important talin-KANK interaction have been elusive.

At a structural level, talin contains an atypical N-terminal FERM domain which is linked with a short linker region to the talin rod region (Fig. 1A)^[4]^. The rod is composed of 13 helical bundles (R1-R13) and a C-terminal dimerization domain (DD) ^[4]^. Whilst the talin FERM domain binds to integrins ^[8]^, the helical bundles in the rod bind to actin and a large number of regulatory proteins ^[9–11]^. The association to filamentous actin (F-actin) can be both direct through two actin-binding sites (ABS2, R4-R8 and ABS3, R13-DD) and indirect through the binding and activation of vinculin which also has an ABS^[12]^. Forces associated with actomyosin activity induce talin conformation changes that can unmask binding sites for vinculin (vinculin binding sites; VBS) and actin (ABS2)^[12, 13]^. We showed previously that constitutively active forms of vinculin that are C-terminally truncated can lock the talin in an activated conformation ^[13, 14]^. When expressed in cells, these active vinculin forms, that contain the talin-binding N-terminal Vd1 domain (or equivalent lacking Vd5, vin880), stabilise FAs even when actomyosin-mediated tension is blocked through inhibitors^[14]^.

**Figure 1.**
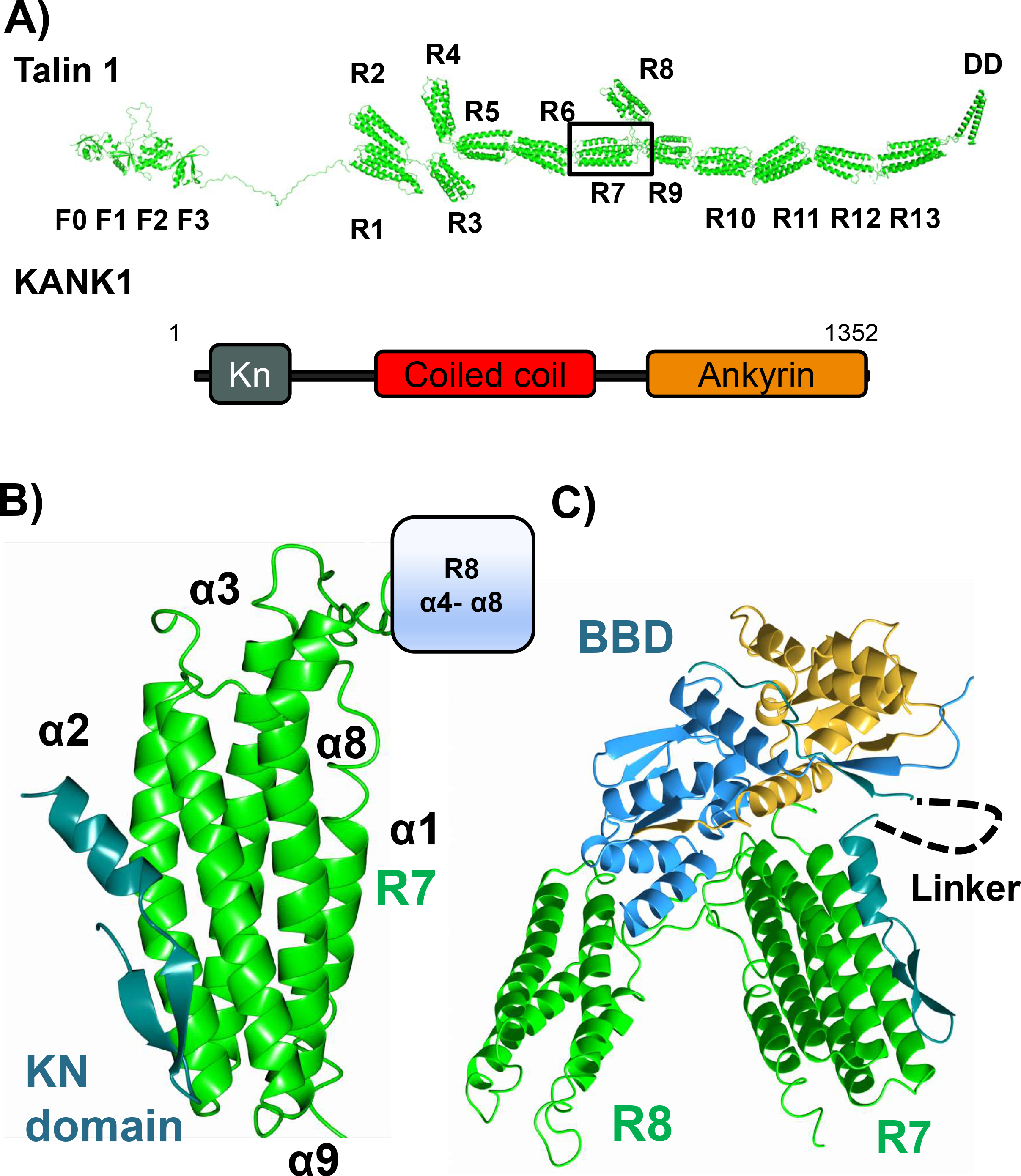
The structure of talin R7 in complex with the KN domain of KANK1. **A)** Talin contains an N-terminal FERM domain connected to a rod region (R1-R13) composed of thirteen 4- and 5-helix bundles. The KANK1 binding site is on the R7 helical bundle. **B)** The KN domain (cyan) binds to talin R7 (green) between helices α2-α9 with no change in any of the helical positions. **C)** The BTB chaperone works by capturing the talin-KANK1 complex in a readily crystallisable lattice.

The talin rod also binds to another class of proteins which bind to the folded rod domains called LD motifs. LD motifs have been identified in multiple talin-binding partners including RIAM^[15]^, DLC1^[16]^, CDK1^[17]^ and paxillin ^[16]^, and are characteristically small amphipathic α-helices (I/LDxØØxØØ consensus sequence where Ø denotes a hydrophobic residue). These LD motifs commonly pack against talin rod domains using a helix addition mechanism and are best exemplified in multiple structures with the talin R8 4-helix bundle ^[15–17]^. KANK (isoforms 1-4) proteins also contain a predicted LD motif in their KN motif region that binds to the 5-helix bundle R7 (Fig. 1A) ^[5, 6]^. However, whereas many of the other binding partners bind to multiple rod domains, KANK binding to talin seems unique since they do not share the promiscuity of binding partners such as paxillin and RIAM. Understanding this novel interaction is therefore important to understand how such specificity for R7 occurs but attempts to crystallise the talin-KANK complexes have been unsuccessful.

In this study, we aimed to gain specific structural detail about the talin-KANK1 interaction and its function in regulating KANK1 localisation to FAs. Using a novel type of crystallographic chaperone, we established a technology that allowed us to solve the structure of an engineered complex that contains the authentic R7-KN interface ^[18]^. Surprisingly, we find that the KN motif is not a conventional LD motif, as instead of being a single helix it instead forms a discrete folded domain that participates in a new type of talin binding interface. Using this new structural information, we designed single-point mutations that disrupt full-length (FL) talin-KANK1 interactions. We show these point mutations completely abolish KANK1 FA localisation, demonstrating that the interaction with talin is essential for KANK1 recruitment to cell-matrix adhesion sites. Stabilising FAs using a constitutively active form of vinculin in parallel with actomyosin inhibitors showed that F-actin directly excludes KANK1 from the core adhesion. Our data lead to a model where actomyosin contractility regulates talin conformation to either promote paxillin-vinculin interactions in the core adhesion or promote KANK1 interactions in the adhesion periphery.

## Results

### Determination of the talin-KANK1 complex using a non-covalent crystallisation chaperone

KANK proteins share a common overall domain structure, each with an N-terminal KN motif responsible for direct interaction with the talin rod domain R7 (Fig. 1A). However, despite the biochemical characterisation of this interaction, the atomic level detail of this interaction was lacking. Therefore, we set out to determine the crystal structure of the talin-KANK1 complex using synthetic KN peptides in complex with recombinant talin R7R8. Attempts to crystallise the complex using standard screening methods including multiple peptide variants, ligand ratio and protein concentrations failed to produce crystalline material and we postulated that the KN motif peptides directly inhibited crystallographic packing.

To overcome this common bottleneck in protein crystallography we generated a new version of Affinity Capture Crystallography (ACC), a method which uses the homodimeric BTB domain of BCL6 as a non-covalent crystallisation chaperone ^[18]^. This and similar approaches use proteins that readily crystallise to donate interfaces and symmetry elements to enable the crystallisation of difficult targets ^[19, 20]^. The procedure requires the expression of a fusion protein containing the monomeric protein of interest with a C-terminal BCL6 Binding Domain (BBD) peptide from its natural binding partner Nuclear Co-repressor 1 (NcoR1). The BBD tag confers a constant high affinity for the BCL6-BTB-homodimer that contains two BBD binding sites (lateral grooves, Fig. S1A) which were modified to be primed for crystal contacts in high-ionic strength conditions. The benefit of this approach is that the chaperone provides an immediate 2-fold symmetry axis that readily packs via multiple potential modes (Fig. S1B). We synthesized a fusion peptide of the mouse KANK1 KN motif (mouse residues 30-60; Fig. S2A) linked to the NcoR1 BBD sequence by a short triglycine linker (KN1_BBD,_ Fig.S2B). A homogenous ternary complex of the BCL6-R7R8-KN_BBD_ was made and purified using size exclusion chromatography (Fig. S2C). The resulting complex was readily crystallised (Fig. S2D) and enabled us to collect X-ray diffraction data of the BCL6-R7R8-KN_BBD_ complex to determine the structure by molecular replacement (Fig. 1B).

The crystal structure revealed how the BTB chaperone supported the crystallisation of the R7-KN motif complex. It shows that the BTB chaperone has donated a back-back interface between BTB homodimers and created a crystallographic tetramer parallel to the *a*-axis (Fig. S3A), and additional contacts were donated from talin R7 (Fig. S3B) that formed a 3-fold homotrimeric complex on the *c*-axis. Overall, the asymmetric unit contained a BCL6 homodimer and a single R7R8 molecule bound to the KN_BBD_ peptide (Fig. 1C). Both the KN region and the NcoR1_BBD_ regions were well resolved in the F_*0*_-F_C_ map, the 2F_*0*_-F*_C_* map and simulated annealing composite omit maps (contoured at 1σ, Fig. S4A) where they shared a similar B-factor distribution of 136.36Å^2^ and 139.54Å^2^, respectively. In the structure, only one of the two lateral grooves of the BTB is occupied due to an unexpected interface between BCL6 and R8 (Fig. S4B) that occludes access to the upper lateral groove sterically restricting corepressor access on a single side. Our strategy defines a new tactic in the determination of challenging protein complex structures and has revealed the structural basis of the talin-KANK1 interaction.

### The KN motif is a novel domain

All previously solved complexes with LD motifs have shown the LD motif to adopt an α-helical conformation (Fig. 2A). In contrast, the KN domain has a novel fold comprised of a β-clasped α-helix. In this fold, the anti-parallel β-clasp is sustained by intramolecular hydrogen bonds between the backbone residues, V33, Q34, T35, P36, F38 and Q39 that connect to and stabilise, the C-terminal α-helical region. Whereas most amphipathic helices tend to have only helical propensity in isolation, this clasp-like structure maintains the rigid three-dimensional epitope with both charged and hydrophobic faces (Fig. S5A). The KN domain defines a new class of talin recognition partners.

**Figure 2.**
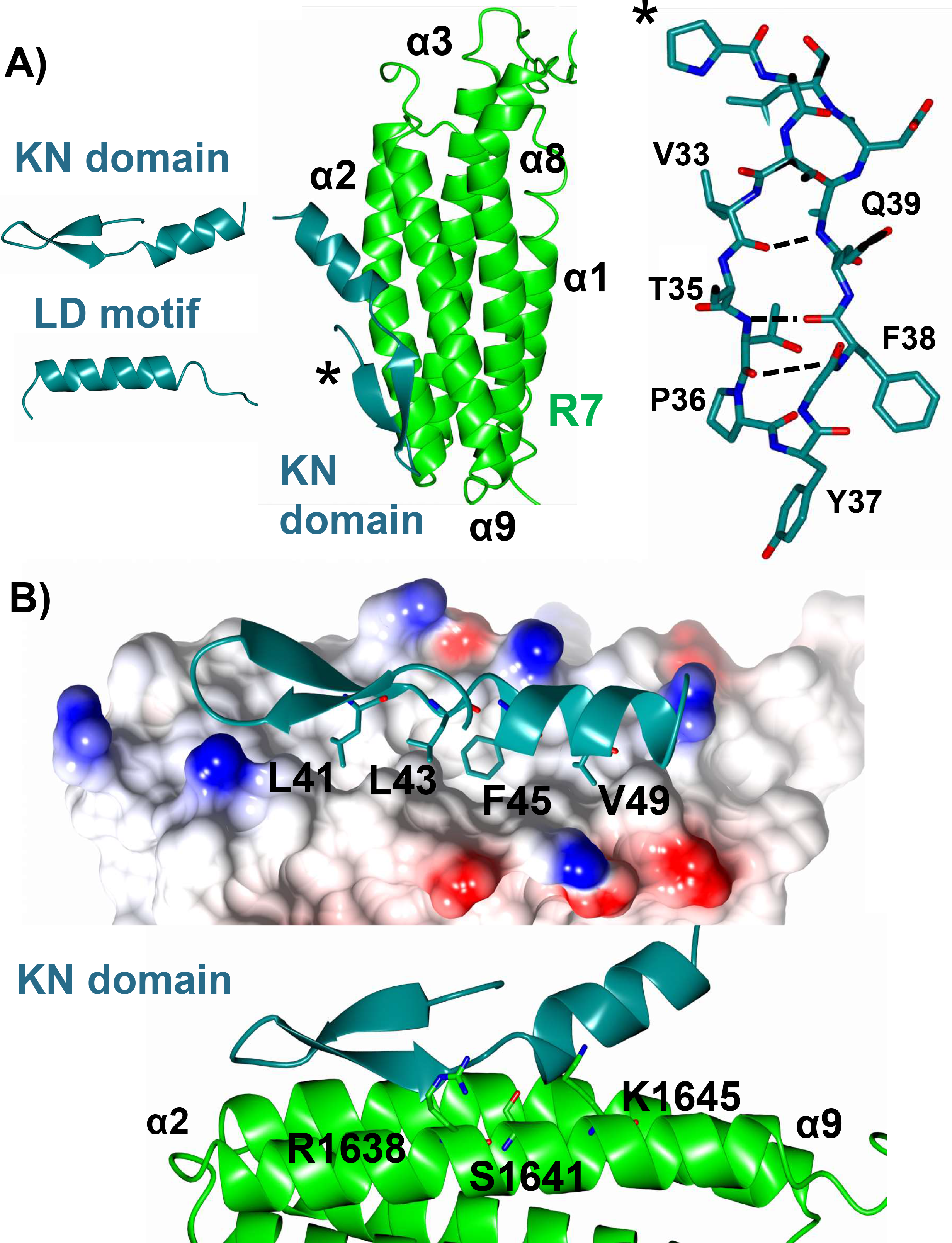
Solution mapping of the talin-KANK1 interface. **A)** The KN domain is a novel fold and is different to the LD motifs. The crystal structure of the talin-KANK1 complex reveals a novel arrangement where intramolecular hydrogen bonds maintain the compact three-dimensional fold of the KN domain. **B)** The anti-parallel β-strand maintains a rigid hydrophobic interface mediated by L41, L43, F45 and V49 sidechains, and with carbonyl side chain bonding donated from talin R1638, S1641 and K1645.

### The structure of the talin-KANK1 complex reveals a novel way to engage a helical bundle

The currently available structures of LD motifs bound to talin show that the interaction is mediated via the helical portion of the LD motif interacting with the helical bundle. In contrast, the R7-KN domain interface is driven principally by the KANK1 β-strand that intercalates with the hydrophobic α2-α9 face of the R7 domain. The interaction involves two main regions, firstly the carbonyl backbone of the KN domain β-strand participates in hydrogen bonding with the sidechains of S1637, R1638, K1645, T1649 and R1652 on R7 (Fig. 2B), and secondly, the KN domain signature “LD” region, ^41^LDLDF^45^, where the sidechains of L41, L43, F45 and V49 occupy complementary hydrophobic cavities that pattern the R7 surface (Fig. 2B). Overall, the novel fold of the KN domain, and the unique interface it makes with talin R7 explain the reason for its stringent specificity and high affinity, in contrast to simpler α-helical LD-motifs such as RIAM and paxillin whose talin rod interactions are multiple. The new structure provided the rationale for the design of structure-based mutations to disrupt the interaction.

### KANK1 point mutants that disrupt the talin interaction abolish KANK1 localisation to FAs

We next explored the effect of charge mutations using Nuclear Magnetic Resonance (NMR). The affinity of the KN domain peptide for R7 is tight, K_d_ 1.2μM (Fig. S5B)) and Heteronuclear Single Quantum Coherence (HSQC) measurements of ^15^N-labelled R7R8 with a 2:1 molar excess of KN domain peptide resulted in large chemical shift changes consistent with this high-affinity interaction (Fig. 3). We next tested variants of the KN domain peptide that were designed to perturb the key contacts identified from the structure. These point mutants introduced single negative charges to replace hydrophobic residues involved in the interface including L41E, L43E, F45E and V49E (Fig. S6). Whilst the wild-type KN domain showed large chemical shift changes, the L41E and F45E mutations produced minimal chemical shift changes demonstrating that the interaction between the KN domain and talin had been abolished. Mutations L43E and V49E were also effective but retained partial, albeit attenuated, interactions.

**Figure 3.**
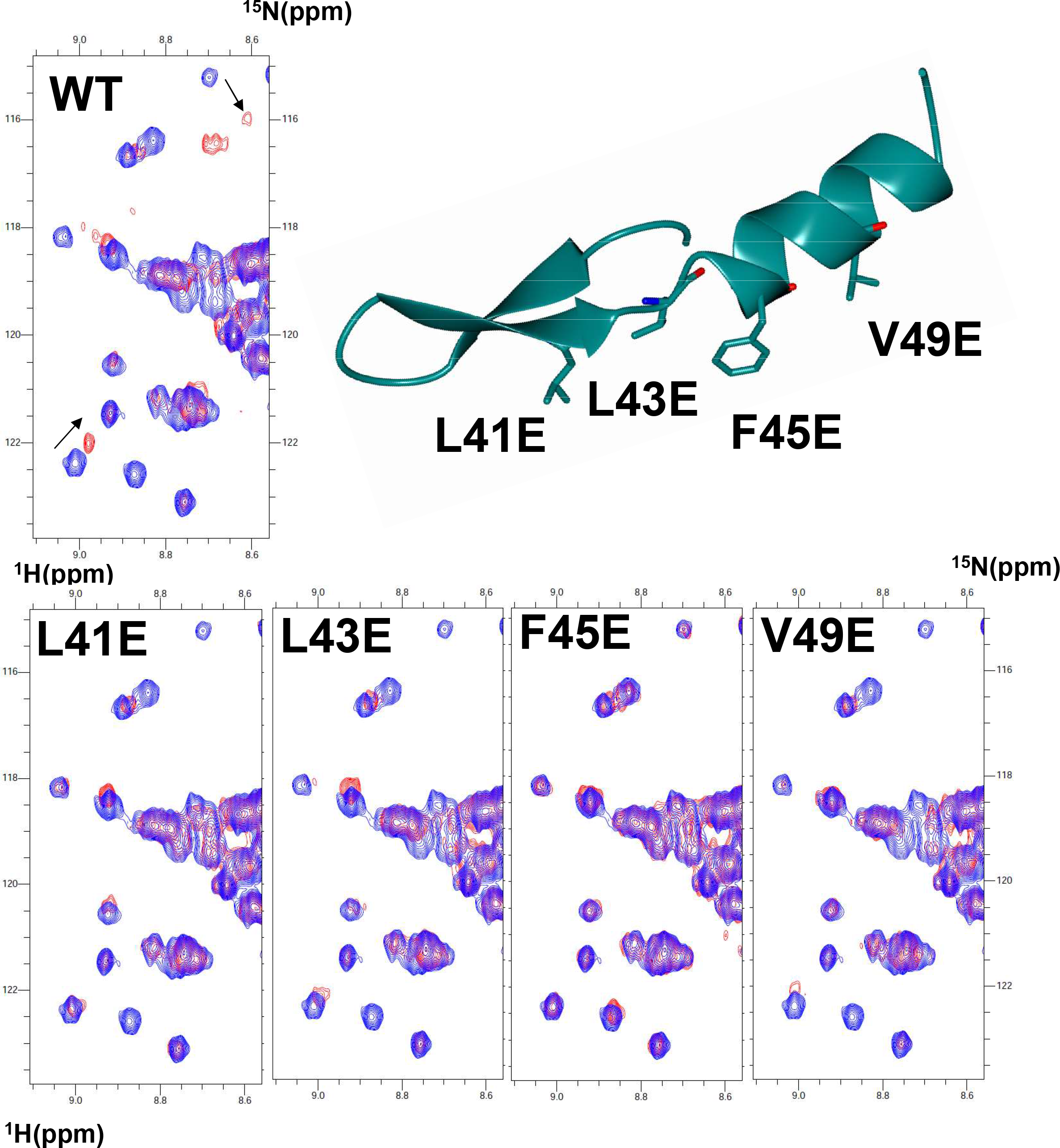
HSQC mapping of the talin-KANK1 interface. ^1^H,^15^N HSQC spectra of 400 μM R7R8 (blue) titrated with a 2-1 molar excess of synthetic KANK1 WT peptide (red) (top). The locations of L41E, L43E, F45E and V49E mutations are shown on the cartoon of the KN domain (right). Bottom: Spectra of R7R8 on own (blue) and in the presence of KANK1 peptides (red).

To examine the effect of these KANK1 mutations on talin interactions in cells, we next used a mitochondrial targeting system (MTS) which we previously used to screen for defined protein-protein interactions^[21]^. In this assay, one protein is fused to a small peptide sequence from the pro-apoptotic protein BAK (cBAK) which leads to its insertion into the outer membrane of mitochondria. Proteins that interact with the cBAK-tagged protein will get recruited to mitochondria, and this recruitment can be verified either by visualising colocalization using fluorescence microscopy (Fig. 4A) or by purification of mitochondria followed by detection of co-precipitates using biochemistry (Fig. 4B, 4C). In such experiments, GFP-talin1-cBAK readily recruits mCherry-KANK1 wild-type (WT) to the mitochondria of both NIH3T3 fibroblasts and HEK293T cells. In contrast, each of the single mutations (L41E, L43E, F45E and V49E) when inserted into KANK1, abolishes GFP-talin-cBAK mediated recruitment (Fig. Fig.S7). To confirm that the interaction was meditated by the R7 domain, we performed the colocalisation assays with truncated talin constructs. Whilst a construct including a rod region starting from R7 to the dimerization motif (GFP-talin1-R7-DD-cBAK) readily colocalised with KANK1, a further truncation comprising R9 to DD (R9-DD) completely abolished colocalisation (Fig. S8A). Moreover, the introduction of the R7 G1404L mutation known to prevent KANK1 association with talin (GFP-talin1G1404L-cBAK) also prevented colocalization (Fig.S8B^[6]^). These findings demonstrate that the talin-KANK1 interaction is sensitive to disruption by single-point mutations.

**Figure 4.**
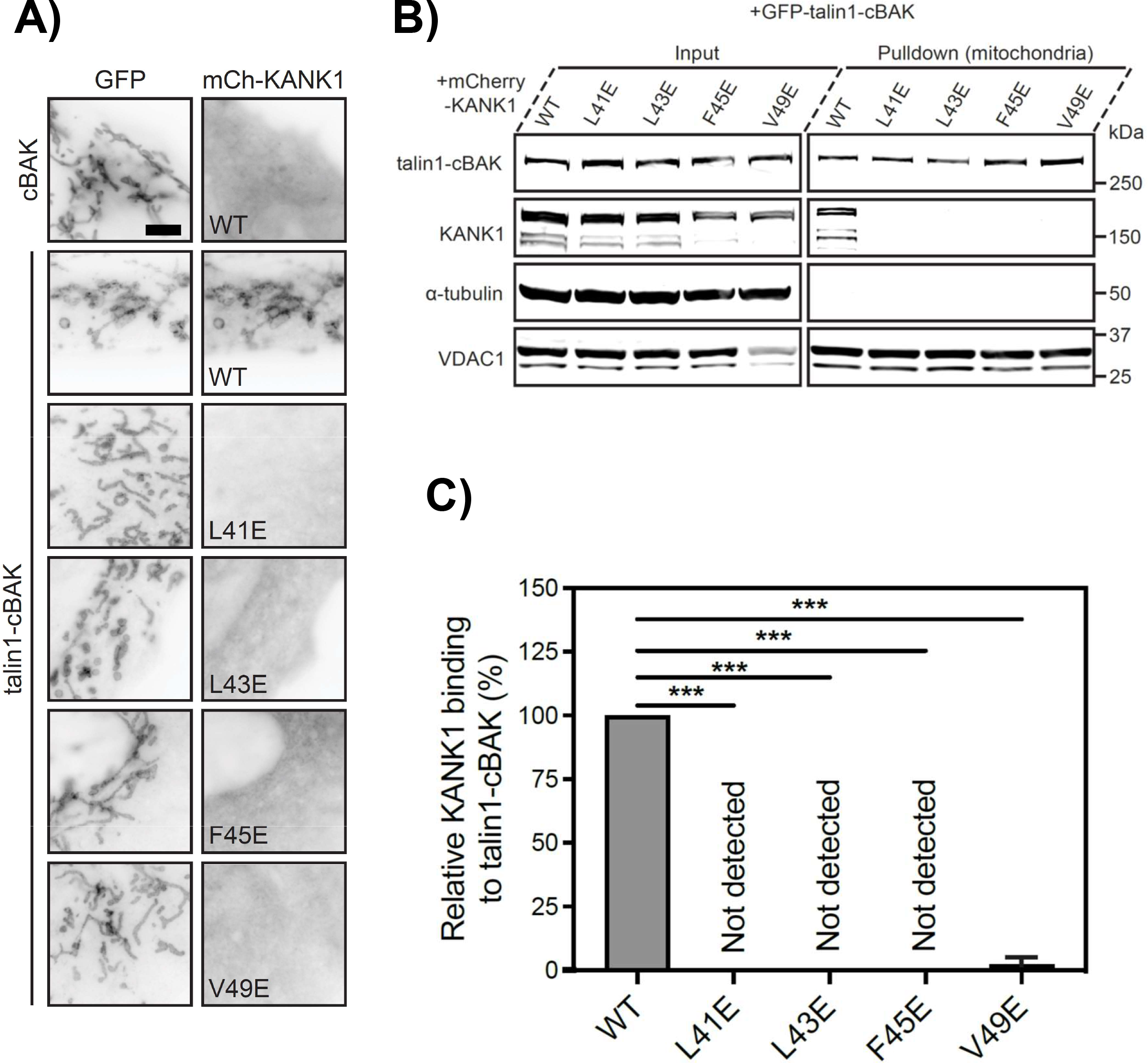
MTS assay and biochemical quantification of talin-KANK1 interactions in cells. **A)** Co-expression of GFP- and GFP-talin-cBAK with mCherry-KANK1 WT, L41E, L43E, F45E and V49E in NIH3T3 fibroblasts, respectively. Note that all mutations abolish the mitochondrial recruitment of mCherry-KANK1. Scale bar 5 μm. **B)** Mitochondria pulldown of KANK1 WT or point mutations in HEK293T cells. **C)** Quantification of KANK1 mitochondrial pulldown from triplicate experiments. Data are normalized to WT. Error bar is SD. *** indicates p<0.001 (Ordinary one-way ANOVA with Dunnett’s multiple comparison test).

### KANK1 is enriched in talin-positive areas of the adhesion belt

To examine KANK1 localisation to FAs we expressed mCherry-KANK1 together with GFP-paxillin in NIH3T3 cells. Whilst both proteins localise to FAs, central areas of FAs that were strongly positive for paxillin were low in KANK1. Reciprocally, KANK1 localised predominantly to the typical “belt-like” structure surrounding a paxillin-enriched central part of FAs and KANK1-enriched areas at the periphery of FAs were low in paxillin. Next, we visualised GFP-talin1 and mCherry-KANK1 in talin knock-out cells which enabled us to observe talin present in the centre of FAs overlapping with paxillin but also in the periphery overlapping with KANK1 (Fig. 5A, Fig. S8C). Co-staining for F-actin showed that actin stress fibres ended in the core of FAs but not in the peripheral KANK1-positive areas (Fig. 5B). To determine the relative effects of our KANK1 talin-binding mutations on KANK1 localisation to FA, we co-expressed WT or mutant mCherry-KANK1 constructs together with GFP-paxillin in NIH3T3 fibroblasts. In these experiments, KANK1 WT readily localised to FAs, and predominantly to the typical “belt-like” structure surrounding the paxillin-enriched central part of FAs. In contrast, all of the KANK1 mutations (L41E, L43E, F45E and V49E) abolished localisation to FAs and the FA-belt (Fig. 5C). Residual mutant KANK1 proteins still localised to regions outside of FAs that were negative for talin but this was likely due to dimerisation with endogenous KANK proteins. The efficiency of our point mutations in abolishing KANK1 localisation to adhesion structures demonstrates that KANK1 localisation to FAs solely depends on its interaction with talin. Moreover, the overlapping distribution of KANK1 with talin in the adhesion belt but the absence of KANK1 from the FA centre that connects with actomyosin suggested the possibility of two populations of talin. One that links to actin, which abolishes KANK1 binding, and a second one which localises to the FA periphery that is devoid of actin.

**Figure 5.**
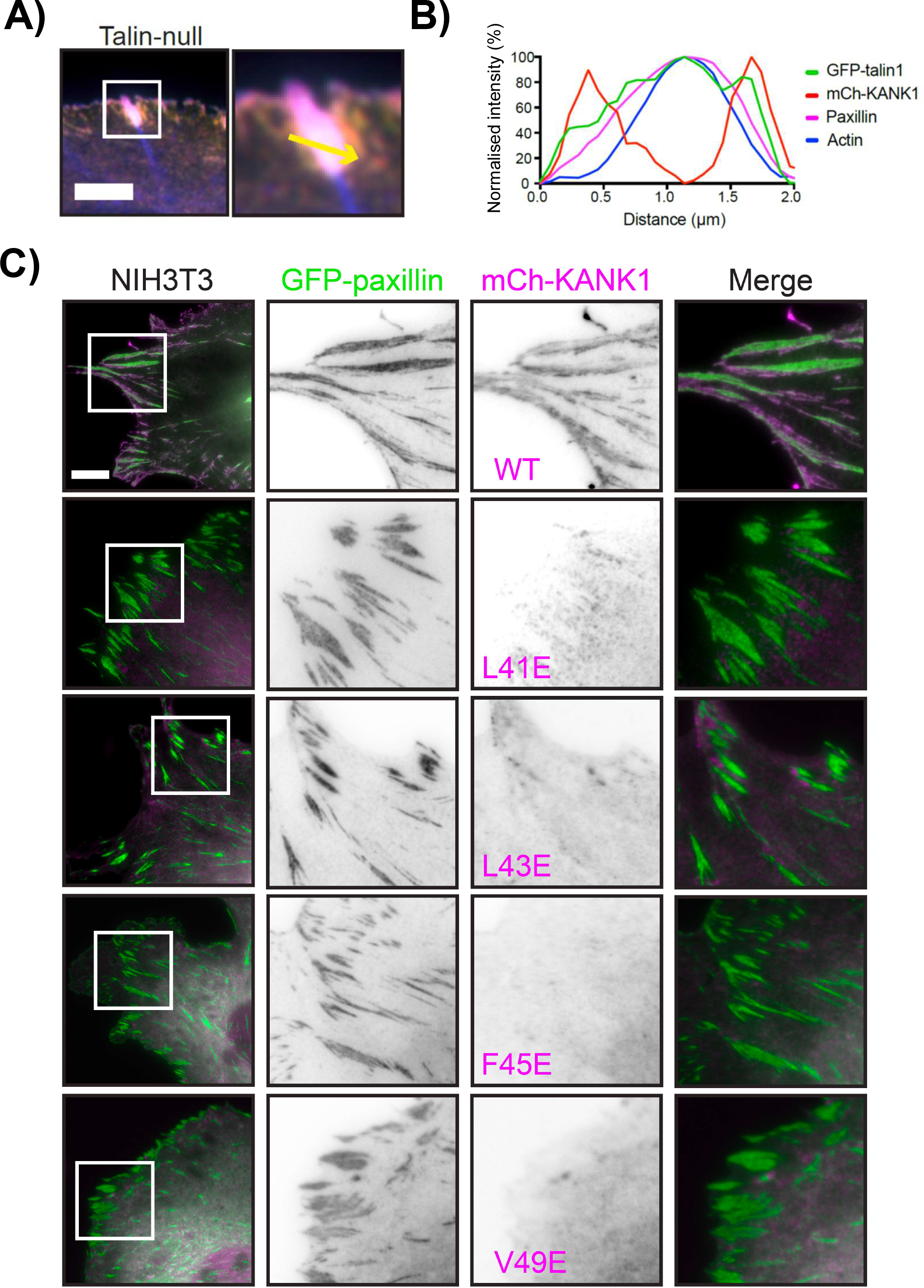
Point mutations in KANK1 abolish adhesion localisation. **A)** Talin null cells were transfected with GFP-talin1 (green), mCherry-KANK1 (red), and immunostained against paxillin (magenta) and actin (phalloidin, blue). Scale bar 5 μm. **B)** Line profile (shown by the yellow arrow in A) indicates normalized fluorescence intensity levels of proteins from a FA in A. **C)** NIH3T3 fibroblasts expressing GFP-paxillin and mCherry-KANK1 wildtype and point mutations L41E, L43E, F45E and V49E. Scale bar 10 μm.

### Dissecting mechanosensitive contributions to KANK1 localisation

Previous reports have shown that forces on talin unmask VBS in the talin rod ^[13]^. We, therefore, hypothesized that actomyosin-induced tension might unfold R7 to expose the VBS in R7 and binding to vinculin would compete with KANK1 at the central parts of FAs. However, experiments in vinculin null MEFs showed similar proportions of KANK1 localising to adhesion belts as in control cells (Fig. S9A). As vinculin does not affect KANK1 localisation, we next examined whether actomyosin forces themselves have a direct impact in triggering the shift of KANK1 from the actin-rich FA centre to the periphery. To explore this possibility we expressed a tailless, constitutively active vinculin construct, vin880, together with KANK1 in NIH3T3 fibroblasts and co-stained these cells for paxillin and actin. Vin880 has been previously shown to produce dramatically enlarged adhesions by maintaining the talin-integrin complex in an activated state which maintains stable FAs even in the presence of inhibitors that block actomyosin-mediated tension ^[14]^. As shown in Fig. 6A, in the control group vin880 produced enlarged adhesions with about 20% adhesions linked to stress fibres showing KANK1 localisation to the FA belt. In contrast, in cells expressing vin880 treated with ROCK inhibitor Y-27632, KANK1 colocalised with paxillin in FAs throughout the whole FA structure (Fig. 6A-6B). Measurement of paxillin/KANK1 colocalisation using the Pearson’s correlation coefficient showed this difference between control and Y-27632 treated group was significant (Fig. 6C-6D, Fig. S10). These data demonstrate that actomyosin-mediated tension modulates KANK1 localisation to FAs with increased actomyosin contractility preventing KANK1 localisation to the central part of adhesions.

**Figure 6.**
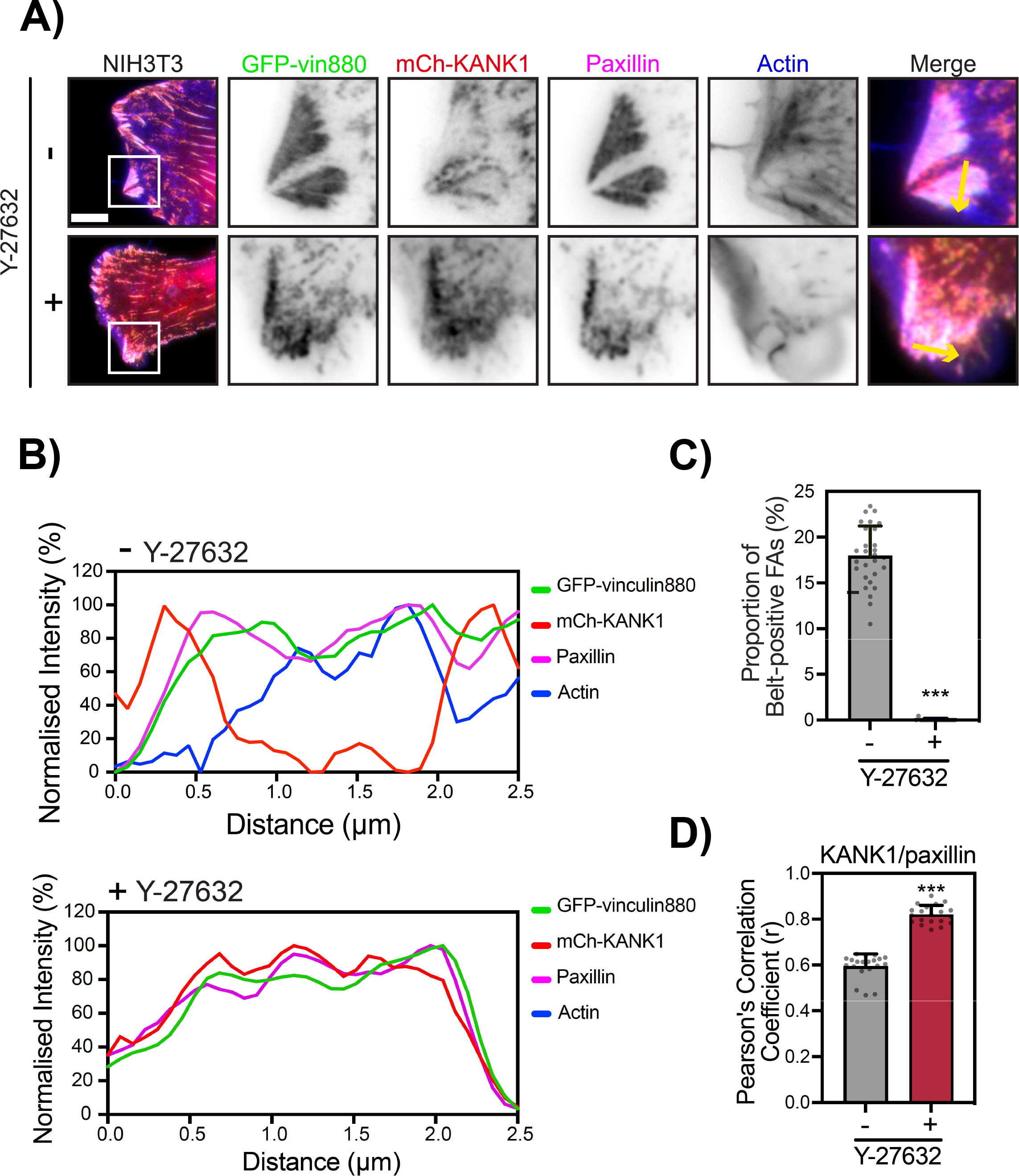
Actomyosin controls the localisation of talin-KANK1. **A)** NIH3T3 fibroblasts expressing GFP-vin880, mCherry-KANK1, and immunostained against paxillin and actin. Cells were treated for 60 minutes with Y27632 (+) or water (−) with associated line profiles shown in **B**. Scale bar 10 μm. **C)** Percentage of belt-positive FAs in Y27632 (+) or water (−) treated cells. FAs with a size over 0.3 μm^2^ were counted. -:17.90±3.27% (n=28); +: 0.02±0.10% (n=17). **D)** Pearson’s mean correlation coefficient of KANK1/paxillin overlap in Y27632 (+) or water (−) treated cells. 20 individual images (40*40 μm) of FA area from each group were measurement. -: r=0.59±0.05; +: r=0.82±0.04. Error bars are SD. *** indicates p<0.001 (Welch’s t test).

### The talin-KANK1 connection organises the assembly of the CMSC

As KANK proteins are central components of the CMSC, we next sought to determine the effect of our F45E KANK1 mutation on the localisation of α/β liprin proteins. In line with previous findings our analysis of NIH3T3 cells showed that both α/β liprins-KANK1 decorate both the cellular cortex and the adhesion belt ^[6, 7]^ (Fig. 7). Introduction of KANK1 F45E mutation showed loss of α/β liprin from both the adhesion belt and the cellular cortex. These findings demonstrate that the talin-KANK1 connection is vital for the ordered assembly of the CMSC both around and recruited to adhesions.

**Figure 7.**
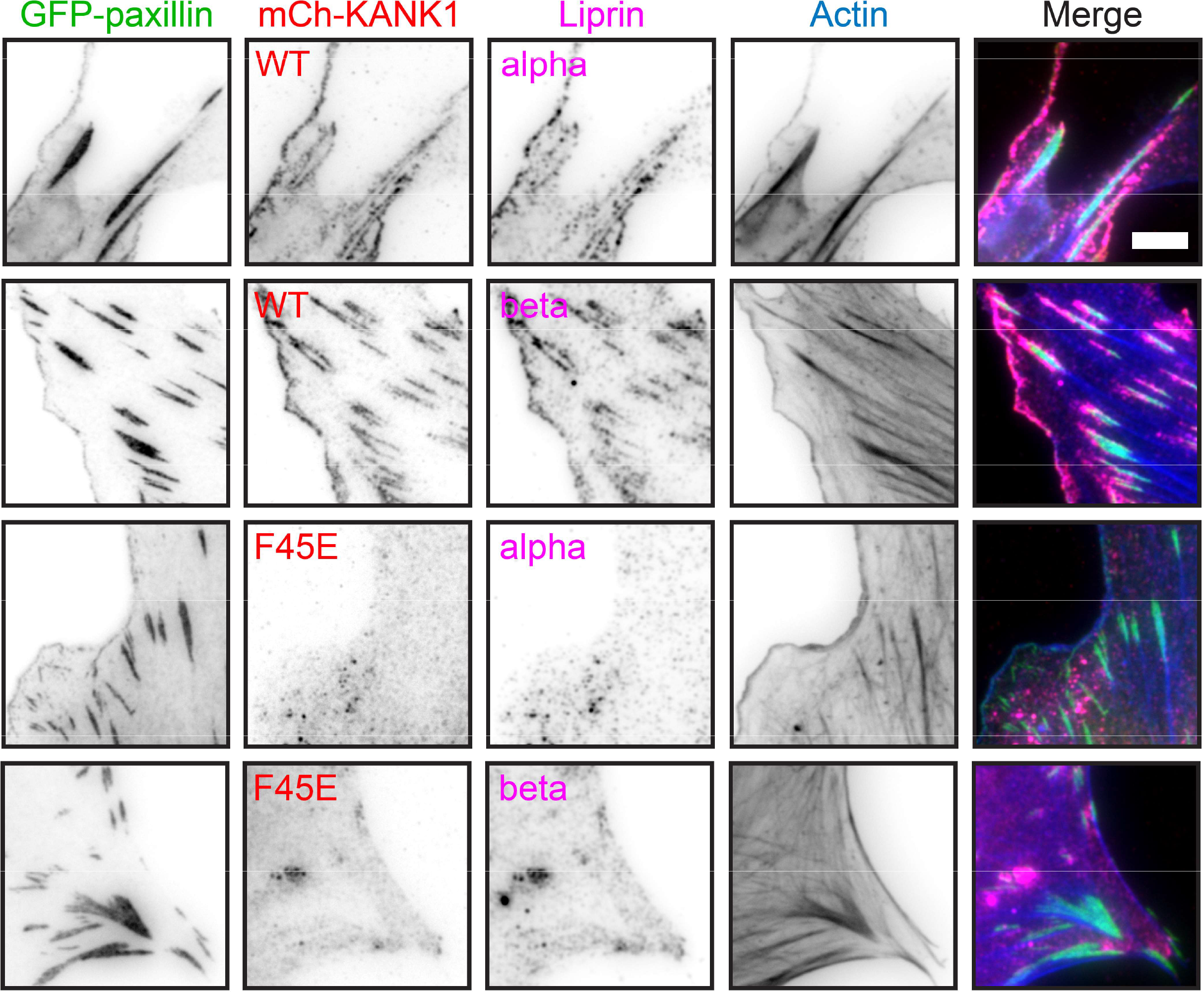
The Talin-KANK1 connection controls CMSC assembly. NIH3T3 fibroblasts were co-transfected with GFP-paxillin and either mCherry-KANK1 WT or F45E, and immunostained against either α or β liprin (magenta) or F-actin (phalloidin, blue). Scale bar 5 μm.

## Discussion

The interaction of talin-KANK1 is critically involved in the recruitment of microtubules to focal adhesions ^[5, 6]^. Yet structural details of this important complex have been elusive. In this study, we developed a new form of crystallography approach to determine the structure of the talin-KANK1 complex. This structure revealed that the KN motif has a novel folded structure and is a KN domain which binds the talin R7 domain with high specificity. Our data enabled us to design single-point mutations that abolish the interaction. We confirmed that KANK1 binding to talin is essential for its localisation to FAs but also demonstrated that talin engagement with actomyosin prevents KANK1 binding to talin.

Determining the structure of the talin-KANK1 complex relied on the development of a new crystallographic approach that allowed us to resolve authentic binary protein-protein complexes. The use of this method enabled us to map the R7-KN domain interface and crucially revealed that the KN motif represents a compact novel protein fold comprising a β-clasp that stabilises an α-helical region. There are nine 5-helix bundles in the talin rod of which only R7 and R11 have so far been shown to bind LD motifs. RIAM binds R11 through a conserved charge-charge interaction facilitating the helical packing of an LD motif against the helical bundle similar to how it engages the 4-helix bundles ^[16]^. R7 does not bind KANK1 in this way, instead, the β-clasp of the KN domain engages a hydrophobic groove localised between α2-α9 on the R7 surface, with a three-dimensional epitope containing the signature ^41^LDLDF^45^ sequence. This β-clasp forms a helical cap, stabilising the helical conformation of the C-terminal region and enhancing its affinity and specificity for R7. Interestingly, this sequence also forms part of the predicted KANK nuclear export sequence (NES), the evolutionary progenitors of the LD motif from whom they diverged 800 million years ago^[22, 23]^. It will be interesting to understand in future studies how the KN domain and LD motif have diverged in evolutionary terms to engage talin by independent binding modes.

Fluorescence polarization measurements confirmed previous observations that the KN domain is the highest affinity talin rod binder identified to date. The crystal structure enabled precise point mutants to be designed that disrupt the interaction that we validated using 2D-NMR *in vitro* and a mitochondrial targeting assay in cells. The fact that these mutations prevent KANK1 localisation to FAs demonstrated that the KANK1 interaction with talin is essential for its recruitment to cell adhesion sites. This observation is in line with previous reports showing that mutations in talin R7 similarly abolish KANK recruitment to FAs ^[6]^, but rules out the hypothesis that the disordered, coiled coils or ankyrin repeat regions may help retain it in adhesion structures ^[5, 6]^.

In previous studies, the differential distribution of talin and KANK proteins in adhesion sites left speculation about the contribution of other proteins in KANK binding at adhesions. However, a more detailed analysis of the fine distribution of talin and KANK1 showed that talin is also present in the, sometimes rather striking, belt area that KANK1 occupies around FAs. Our results show that the area of FAs highest enriched for F-actin is lowest for KANK1 and vice versa. Previous studies already forwarded the hypothesis that actomyosin negatively impacts KANK localisation to adhesions ^[5, 6]^. One study found that *in vitro* F-actin and KANK2 compete for the ABS2 region explaining reduced force transmission integrin-mediated adhesions^[5]^, and another observed that KANK gradually occupied the remaining adhesions upon actomyosin inhibition^[6]^. This led us to explore the relation between the mechanical state of talin, KANK1 localisation and the relative contribution of vinculin and F-actin.

Vin880, a constitutively active vinculin construct, maintains focal adhesion structures even in the presence of ROCK1 inhibitors but abolishes the conformational changes of actomyosin-induced tension. Our data demonstrated that in the absence of F-actin KANK1 will colocalise perfectly with paxillin and vin880. Mechanistically, and in line with our finding in vinculin null cells, our data rule out the mechanical recruitment of vinculin as a driver of KANK1 adhesion exclusion and highlights the function of F-actin. It also demonstrates that the R7R8 double domain module in talin is folded in these experiments and able to participate in protein interactions that would be thought to be mutually exclusive with vinculin binding. Previous reports have shown that increasing the hydrophobic core of domains, such as R3^[4]^, can stabilise the fold. Therefore it may be possible that the hydrophobic interface of the R7-KN domain increases the mechanical stability of R7 to permit recruitment and inhibit the talin-vinculin association^[24]^.

KANK1 proteins are part of the CMSC that decorate the leading edge and cellular cortex as well as the adhesion belt ^[6, 7]^. Therefore, we sought to examine the localisation of α/β liprin proteins in response to our F45E mutation. Our data demonstrated that this mutation abolished both adhesion localisation and also strikingly α/β liprin organisation at the lamellipodia. Our data highlight the reciprocal cross-talk between FA and the CMSC and how talin maintains both assemblies. Overall, our findings have advanced crystallography and provided atomic-level detail about the elusive talin-KANK interaction. In future, this insight will facilitate the design of small molecular inhibitors to disrupt the talin-KANK1 axis, which given the importance of KANK proteins in disease, will enable precise dissection of this important linkage.

## Acknowledgements

X-ray data were collected as part of the University of Manchester BAG allocation. We wish to thank the Bioimaging facility at the University of Manchester (UoM) and the NMR centre for structural biology at the University of Liverpool. The Zacharchenko and the Ballestrem laboratories are part of the Wellcome Trust Centre for Cell-Matrix Research, University of Manchester, which is supported by core funding from the Wellcome Trust (grant number 203128/Z/16/Z). CB acknowledges BBSRC for funding (BB/V016326/1). The Bioimaging Facility microscopes were purchased with grants from the BBSRC, the Wellcome Trust, and the University of Manchester. We also thank Professor Tim Hardingham (UoM) for the critical reading of the manuscript. TZ is funded by the Presidential Research Fellowship, University of Manchester.

## Author contributions

TZ conceived the project and performed biochemistry, X-Ray crystallography and data analysis. CB conceived all cell biology experiments and XL performed these experiments. TZ wrote the manuscript with contributions from CB, BTG and XL.

## Materials and Methods

### Protein expression and purification

Mouse talin-1 (P26039) R7R8 was expressed and purified as described previously^[9]^. The BCL6 chaperone was expressed and purified as described previously^[18]^. Constructs were verified independently by sequencing.

### Synthetic peptides

Peptides were purchased from GLBiochem (Shanghai). Peptides include the KN1_BBD_ fusion ^1^PYFVETPYGFQLDLDFVKYVDDIQKGNTIKKGGGGITTIKEMGRSIHEIPR^51^ and the KANK1 KN domain (UniProt E9Q238) ^30^PYFVETPYGFQLDLDFVKYVDDIQKGNTIKKC^60^ (for FP), and^30^PYFVETPYGFQLDLDFVKYVDDIQKGNTIKK^60^ (for NMR). The following peptides were used for NMR screening L41E ^30^PYFVETPYGFQEDLDFVKYVDDIQKGNTIKK^60^, L43E ^30^PYFVETPYGFQLDEDFVKYVDDIQKGNTIKK^60,^ F45E ^30^PYFVETPYGFQLDLDEVKYVDDIQKGNTIKK^60^ and V49E ^30^PYFVETPYGFQLDLDFVKYEDDIQKGNTIKKC^60^.

### Fluorescence Polarisation Assay

For determination of the WT-KANK1 binding constant the BODIPY-TMR coupled peptides dissolved in PBS (137 mM NaCl, 27 mM KCl, 100 mM Na_2_HPO_4_, 18 mM KH_2_PO_4_, pH 7.4), 5 mM TCEP, and 0.05% (v/v) Triton X-100 were used at a final concentration of 0.5 μM. Uncoupled dye was removed using a PD-10 gel filtration column (GE Healthcare). Fluorescence polarization measurements were recorded on a BMGLabTech CLARIOstar plate reader and analysed using GraphPad Prism (version 6.07). K_d_ values were calculated by nonlinear curve fitting using a one-site total and nonspecific binding model.

### Nuclear Magnetic Resonance

NMR spectra were collected on Bruker Avance III 800 MHz spectrometer equipped with CryoProbe. Experiments were performed at 298 K in 20 mM sodium phosphate (pH 6.5) and 50 mM NaCl, 3mM β-mercaptoethanol with 5% (v/v) ^2^H_2_O.

### X-ray crystallography

Initial sparse matrix crystal approaches using the strategies described previously for R7R8 complexes failed to produce any crystalline material, as did further modulation of protein/peptide concentration. We used a synthetic peptide containing the NCoR1_BBD_ connected via a triglycine linker to the C-terminus of the KANK1 KN domain (residues 30-60) peptide ^1^PYFVETPYGFQLDLDFVKYVDDIQKGNTIKK-GGG-GITTIKEMGRSIHEIPR^51^. This KN1_BBD_ peptide facilitated the formation of a ternary R7R8-BCL6-KN_BBD_ complex with the BCL6 non-covalent chaperone that was readily purified by size-exclusion chromatography. The complex was concentrated to 10 mg/ml and used for crystallographic screening in 20 mM Tris pH 7.4, 150 mM NaCl, and 3 mM β-mercaptoethanol. Crystals were obtained by conventional sparse matrix screening sitting drop vapour diffusion with plates dispensed using a Mosquito Liquid Handling robot (SPT Labtech) with a 1:1 precipitant-precipitate ratio in 400 nl drops. Crystals were obtained in 1 M Ammonium Sulphate, 0.1 M CHES pH 9.5, 0.2 M NaCl, 6% Glycerol and typically after ~3 weeks and before data collection vitrified in mother liquor containing 20% glycerol. Diffraction data were collected on I03 Diamond Light Source using the automated collection mode and integrated using XDS/SCALA, resolution cut-off was determined by CC_1/2_ at 3.4Å (0.289) ^[25, 26]^. Crystals adopted space group H32 and the structure of the complex was solved by molecular replacement using PHASER^[27]^ with the template structure of the BCL6-NCoR1_BBD_ complex (PDB:6XYX). After molecular replacement electron density of both R7, R8 and the KN were visible allowing the placing of R7 and R8 using PHASER, and the unambiguous assignment of the KN domain in COOT^[28]^. Data reduction statistics and refinement information are shown in Table 1 and coordinates and structure factors were deposited to the PDB with the accession code 8AS9.

R_free_ was calculated using 5% of data isolated from the refinement for cross-validation. The highest-resolution shells are shown in parentheses. TLS parameters used chains A, B, C and D.

|  |  |
| --- | --- |
| **Beamline** | **I03 Diamond** |
| Detector | DECTRIS EIGER2 XE 16M |
| Wavelength (Å) | 0.97625 |
| Resolution range (Å) | 61.89-3.40 (3.58-3.4) |
| Space group | R32:H |
| *a*, *b*, *c* (Å) | 207.02 207.02 151.86 |
| $\alpha$, $\beta$, $\gamma$ (°) | 90 90 120 |
| Unique reflections | 17332 (2505) |
| Completeness (%) | 100 (100) |
| Multiplicity | 20.9 (21.8) |
| Mean (I) CC<sub>1/2</sub> | 0.992 (0.289) |
| <I/$\sigma$(I)> | 5.1 (0.5) |
| R<sub>merge</sub> (I) | 0.641 (6.191) |
| R<sub>p.i.m.</sub> (I) | 0.143 (1.356) |
| **Refinement** |  |
| RMSD bonds (Å) | 0.003 |
| RMSD angles (°) | 0.880 |
| R<sub>work</sub>(%) | 24.50 (39.32) |
| R<sub>free</sub>(%) | 28.60 (40.27) |
| **No. of atoms** |  |
| Total | 4704 |
| Macromolecule | 4678 |
| Solvent | 26 |
| **Ramachandran** |  |
| Favoured (%) | 94.43 |
| Allowed (%) | 4.73 |

### Cell lines and transfections

NIH3T3 mouse fibroblasts and HEK293T human epithelial cells were obtained from the American Type Culture Collection (ATCC). The vinculin null and WT mouse embryonic fibroblasts (MEFs) originate from the Eileen Adamson laboratory ^[29]^. All cells were maintained in Dulbecco’s Modified Eagle Medium (DMEM; Sigma) supplemented with 10% fetal bovine serum (FBS, Gibco), 1% L-glutamine (Sigma) and 1% non-essential amino acids (Sigma). Talin1&2 null cells ^[12]^ were cultured in DMEM F-12 (Gibco) supplemented with 10% FBS, 1% L-glutamine, 1% non-essential amino acids and 15□μM HEPES (Sigma). All cells were cultured at 37°C supplied with 5% CO2 and 95% humidity. Transient transfections were performed using Lipofectamine LTX with Plus Reagent (Invitrogen) to NIH3T3 cells, Lipofectamine 2000 (Invitrogen) to talin null cells, and jetPRIME reagent (Polyplus) to MEFs and HEK293T cells, respectively, as per the manufacturer’s instructions.

### Plasmids preparation and site-directed mutagenesis

For the construction of mCherry-KANK1, FL-human KANK1 (generous donation from the Bershadsky lab) cDNA was tagged in the C-terminal site with pmCherry (Clontech) by restriction digestion. Point mutations (L41E, L43E, F45E and V49E) were introduced in mCherry-KANK1 by site-directed mutagenesis (NEB) using the following oligonucleotides: TGGTTATCAAgaAGACTTAGATTTCCTCAAATATG and TAGGGGGTCTCCACAAAG for L41E; TCAACTAGACgaAGATTTCCTCAAATATGTG and TAACCATAGGGGGTCTCC for L43E; AGACTTAGATgaaCTCAAATATGTGGATG and AGTTGATAACCATAGGGG for F45E; CTCAAATATGaGGATGACATACAG and GAAATCTAAGTCTAGTTGATAAC for V49E. Generation of GFP-talin1-cBAK and GFP-cBAK were previously described ^[21]^. G1404L was introduced on GFP-talin1-cBAK by the method above using CAAGGTCCTActtGAGGCCATGACTGG and GAGTTCTCCATGACACTG. To generate GFP-talin1 R7DD-cBAK and GFP-talin1 R9DD-cBAK constructs, R7-DD (3,555 bp) and R9-DD (2,661 bp) were amplified from talin-1 (Mus musculus). Restriction digestion with XhoI and HindIII FastDigest enzymes (Thermo Scientific) was used to linearize GFP-cBAK. DNA assembly was performed to join R7-DD and R9-DD to linearised the GFP-cBAK vector using the NEBuilder HiFi DNA assembly kit (NEB).

### Antibodies and reagents

For fixed cell imaging, cells were cultured in glass-bottom dishes (IBL) coated with bovine fibronectin (Sigma) at a final concentration of 10 μg/ml. Samples were fixed in 4% paraformaldehyde (PFA, Sigma), and warmed to 37°C, for 15 minutes before being washed three times with PBS. For immunofluorescence staining, samples were permeabilised at room temperature with 0.5% Triton X-100 (Sigma) for 5 minutes before being washed three times. The primary antibody rabbit anti-paxillin (clone Y113, ab32084, Abcam) was used at a dilution of 1:200 (in 1% BSA), rabbit anti-liprin alpha (14175-1-AP, Proteintech) was used 1:200, rabbit anti-liprin beta (11492-1-AP, Proteintech) was used 1:200. Secondary antibody Alexa Fluor Plus 647 goat anti-rabbit (Invitrogen) was used at a dilution of 1:500. Actin was visualized using Alexa Fluor Plus 405 Phalloidin (1:500, Invitrogen). Y-27632 (Tocris Bioscience) was diluted in dH20 and used at a final concentration of 50 μM. Before use, the stock was diluted in a pre-warmed medium before being added to cells. Mitochondria isolation from HEK293T cells was performed after 24 hours of cell transfection using the Q proteome Mitochondria Isolation Kit (QIAGEN). Cell lysis and mitochondria homogenisation were conducted as per the manufacturer’s instructions. The purified mitochondrial were stored at −80°C.

### Microscopy

Images of fixed samples in PBS were acquired at room temperature using an Olympus IX83 inverted microscope equipped with a 60x/1.42 PlanApo N oil objective and a QImaging Retiga R6 CCD camera, controlled by Metamorph software. Samples were illuminated using LEDs (UV/Cyan/Green-Yellow/Red, Lumencor) for fluorescence excitation; a Sedat filter set (DAPI/FITC/TRITC/Cy5, Chroma, 89000) was used.

### Protein extraction and Western blot

Protein samples were extracted from cells and mitochondria, respectively, using RIPA lysis buffer (Chromotek) supplemented with protease inhibitors. Protein samples were diluted in LDS sample buffer (4X, Invitrogen) supplemented with sample reducing agent (10X, Invitrogen). Samples were heated at 95°C for 5 minutes before loading on a 4-12% gradient Bis-Tris gel (Invitrogen). MOPS SDS running buffer (Invitrogen) was used and supplied with antioxidants (Invitrogen). The gel was soaked in running buffer and run at 160 V for 75 minutes. The gel was transferred to a 0.45 μm nitrocellulose membrane (Cytiva) and protein at 30 V for 150 minutes, 4°C. The membrane was blocked for 1 hour in 5% skimmed milk (Sigma) in PBS-Tween 20 (0.1%, Sigma). The membrane was probed for anti-GFP (ab183734, abcam), anti-mCherry (1C51, ab125096, abcam), anti-VDAC1 (ab15895, abcam) and anti-α tubulin (DM1α, T6199, Sigma), diluted 1:10000, 1:3000, 1:1500, 1:1500, respectively, in 5% milk (PBS-Tween). Primary antibody signal was detected using goat anti-mouse IgG conjugated to IRDye® 680RD (ab216776, abcam) and goat anti-rabbit IgG conjugated to IRDye® 800CW (ab216773, abcam) secondary antibodies, diluted 1:15000, imaged with an Odyssey CLx imaging system (LI-CO Biosciences).

### Analysis of cell adhesions

FIJI-ImageJ^[30]^ software was used to process all images. Cell-matrix adhesion size was quantified as described previously^[12]^, by subtracting background signal using a rolling ball algorithm, followed by thresholding to select adhesion structures and the Analyze Particles function to quantify adhesions. The line intensity profile of adhesion was generated using the Plot Profile function. The intensity profile was then normalized between 0 and 100% by dividing the plot value by the maximum value and then multiplying by 100. Pearson’s correlation coefficient of fluorescence signals (40 × 40 μm square adhesion area) was measured by subtracting the background signal, followed by automatic thresholding and colocalisation analysis using the Bioimaging and Optics Platform (BIOP) version JACoP plugin^[31]^.

### Graphs and statistical analysis

All graphs and statistical analyses were carried out using Prism 9 (GraphPad). Where appropriate, statistical significance between two individual groups was tested using an unpaired t-test with Welch’s correction. An ordinary one-way analysis of variance (ANOVA) followed by Turkey’s multiple comparison tests was performed to test for significance between tests or more groups. Data distribution was tested for normality using a D’Agostino & Pearson omnibus normality test; a P value >0.05 was used to determine normality. Data are presented as mean ± standard deviation (SD). A P value of 0.05 or below was considered statistically significant. * p<0.05, ** p<0.01 and *** p<0.001.

## Figure legends

**Figure S1.**
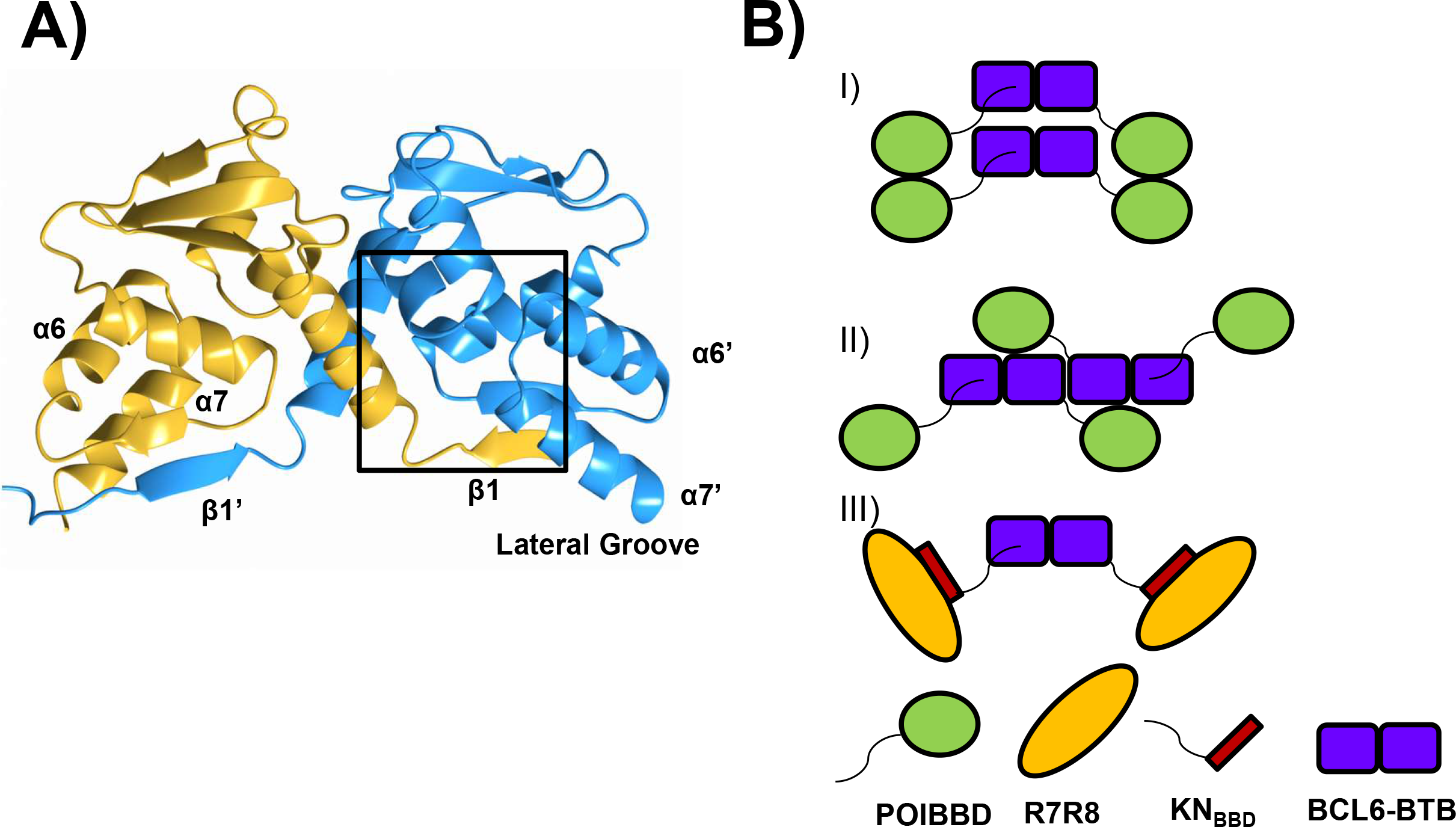
**A)** The BCL6-BTB domain forms a symmetrical, strand exchanged homodimer that binds to two BBD peptides via the lateral groove. **B)** Potential modes of BCL6-BTB Packing (blue), model i) as demonstrated in PDB (PDB:6Y17 with Protein of Interest (POI) connected to a BBD domain (green), **ii)** Hypothesised packing and **iii)** Concept of affinity capture crystallography (ACC) in this work, where a POI is connected to the chaperone with a linkage peptide.

**Figure S2.**
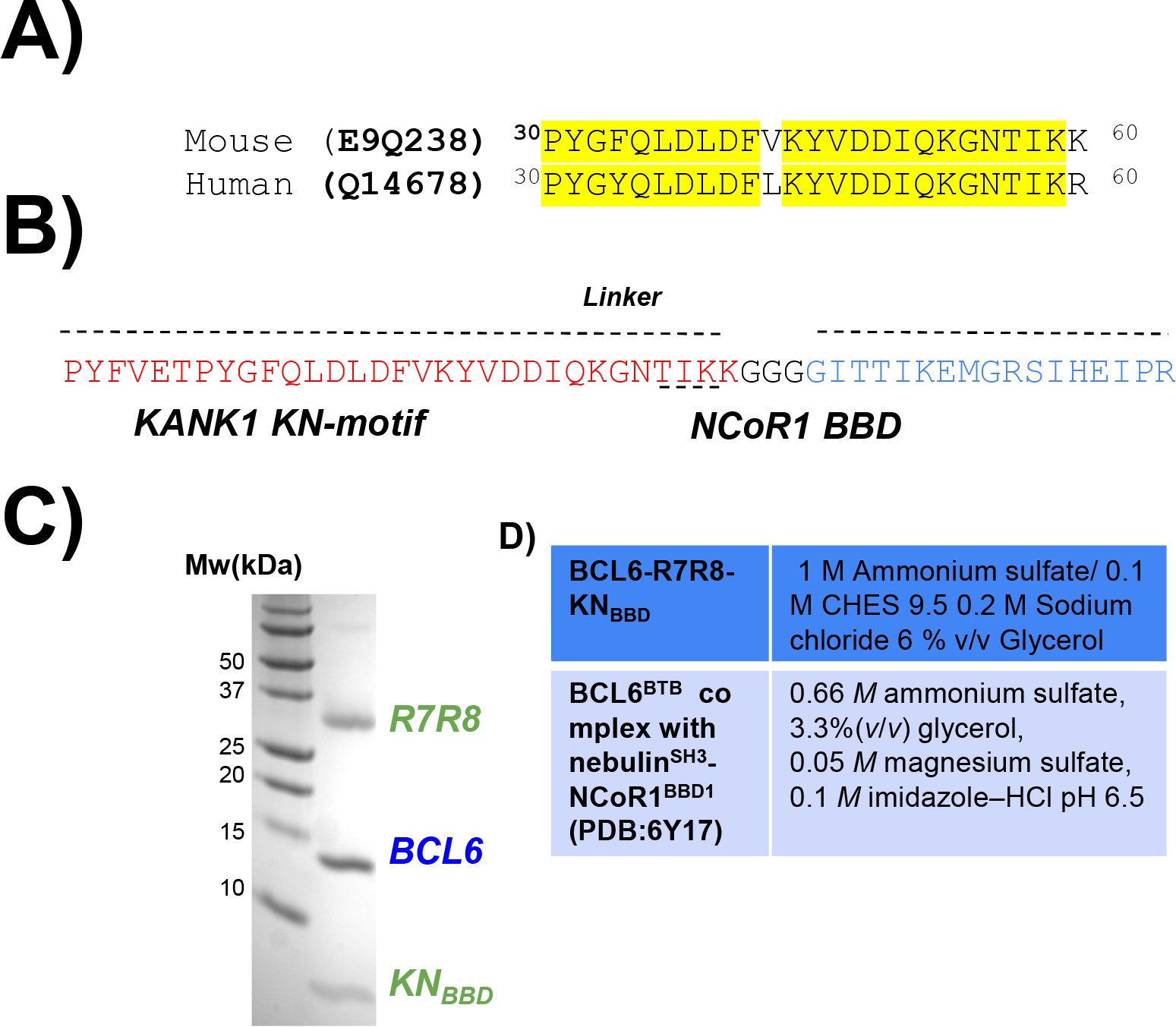
**A)** Sequence alignment of the mouse and human KN domains. **B)** Sequence of the KN_BBD_ peptide with mouse KANK1 KN-motif shown in red, connected to the NCoR1 BBD sequence in blue by a triglycine linker. **C)** 16% Tris-Tricine SDS-PAGE gel of the eluted maxima from a Superdex 200 10/300 increase column**. D)** Table of conditions between of the BCL6-R7R8-KN_BBD_ and the BCL6^BTB^ complex with nebulin^SH3^-NCoR1^BBD1^ (PDB:6Y17).

**Figure S3.**
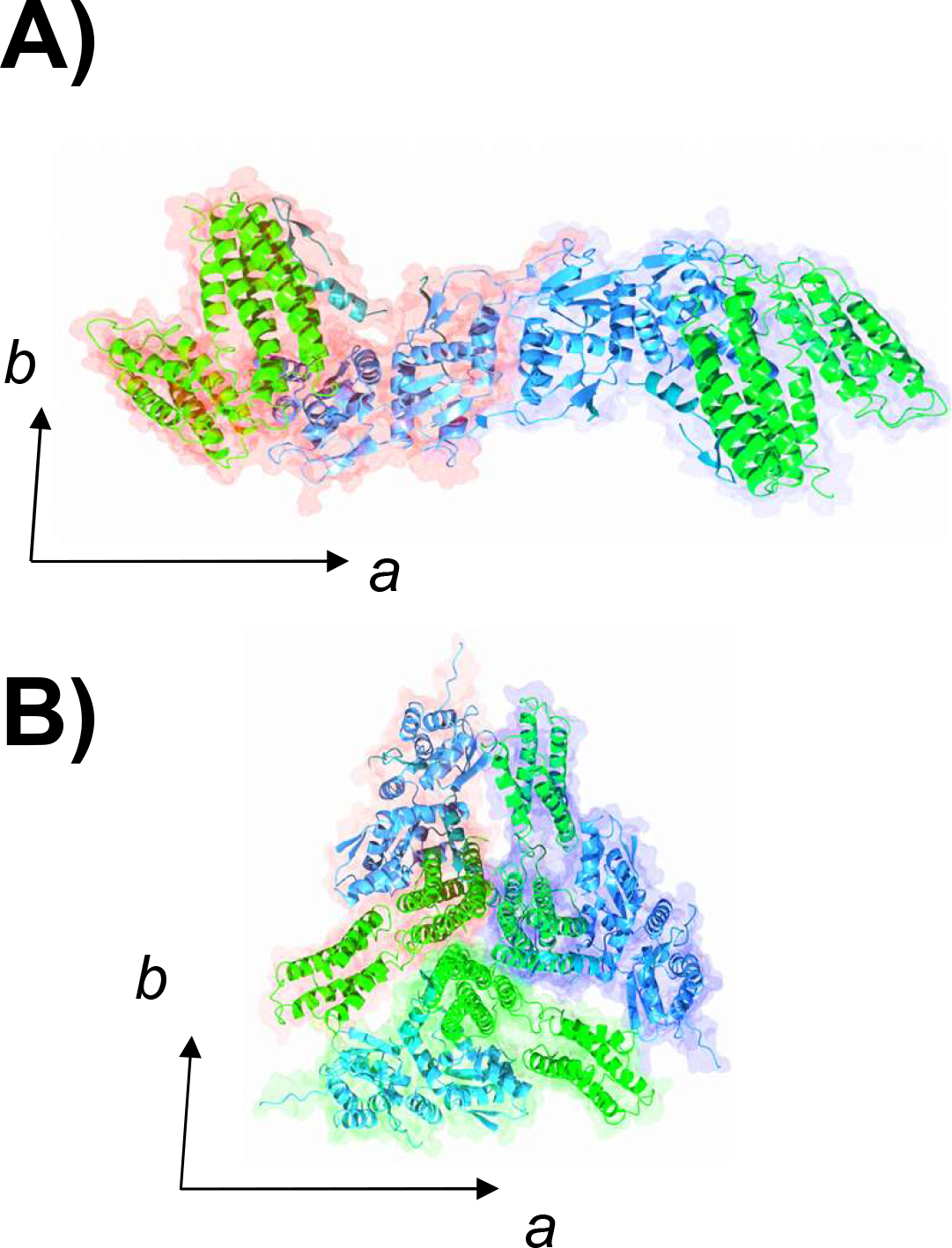
Analysis of the contribution of the chaperone to crystallographic packing. **A)** Assembly parallel the a-axis**. B)** View down the c-axis.

**Figure S4.**
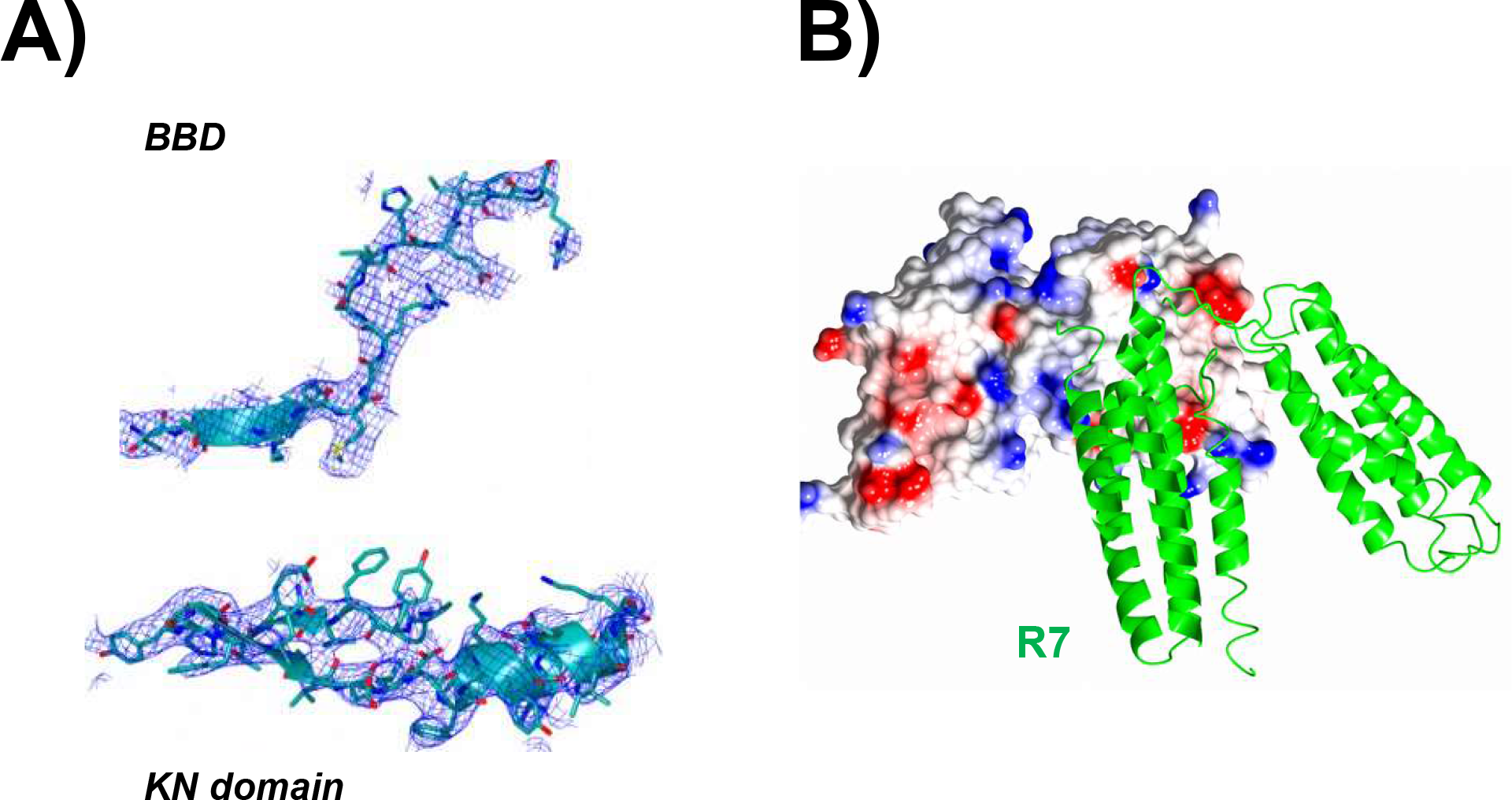
**A)** 2F_0_-F_C_ simulated annealing composite omit map of the BBD peptide and KN domain. **B)** Poisson-Boltzmann Distribution of BCL6 in a tethered complex with R7R8.

**Figure S5.**
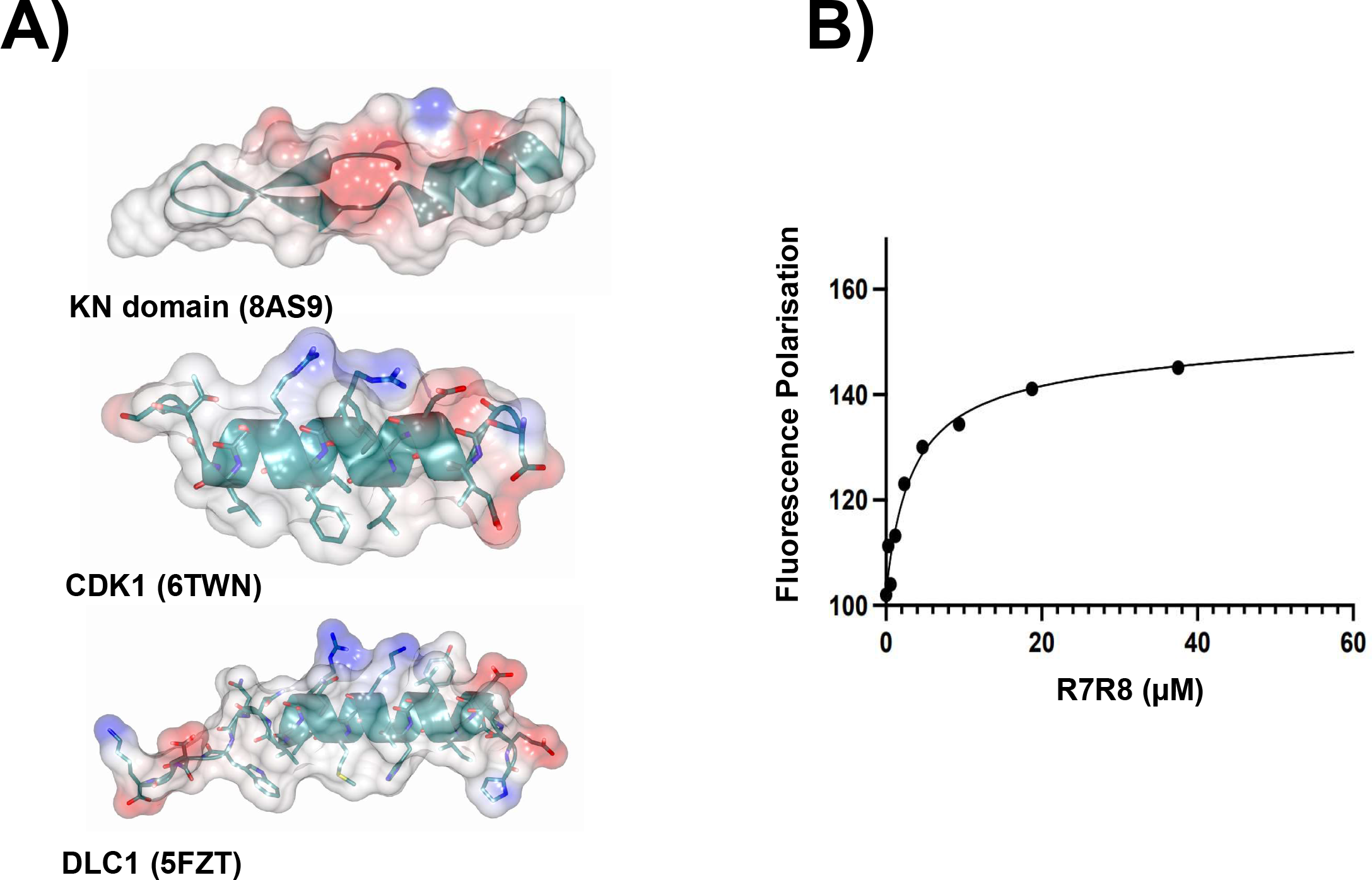
**A)** Poisson-Boltzmann distribution maps of the KANK1 KN domain and the amphipathic talin-binding LD motifs from CDK1 and DLC1. **B)** Fluorescence Polarisation measurements show that the affinity between R7 and KN domain peptide has a K_d_ of 1.2 μM.

**Figure S6.**
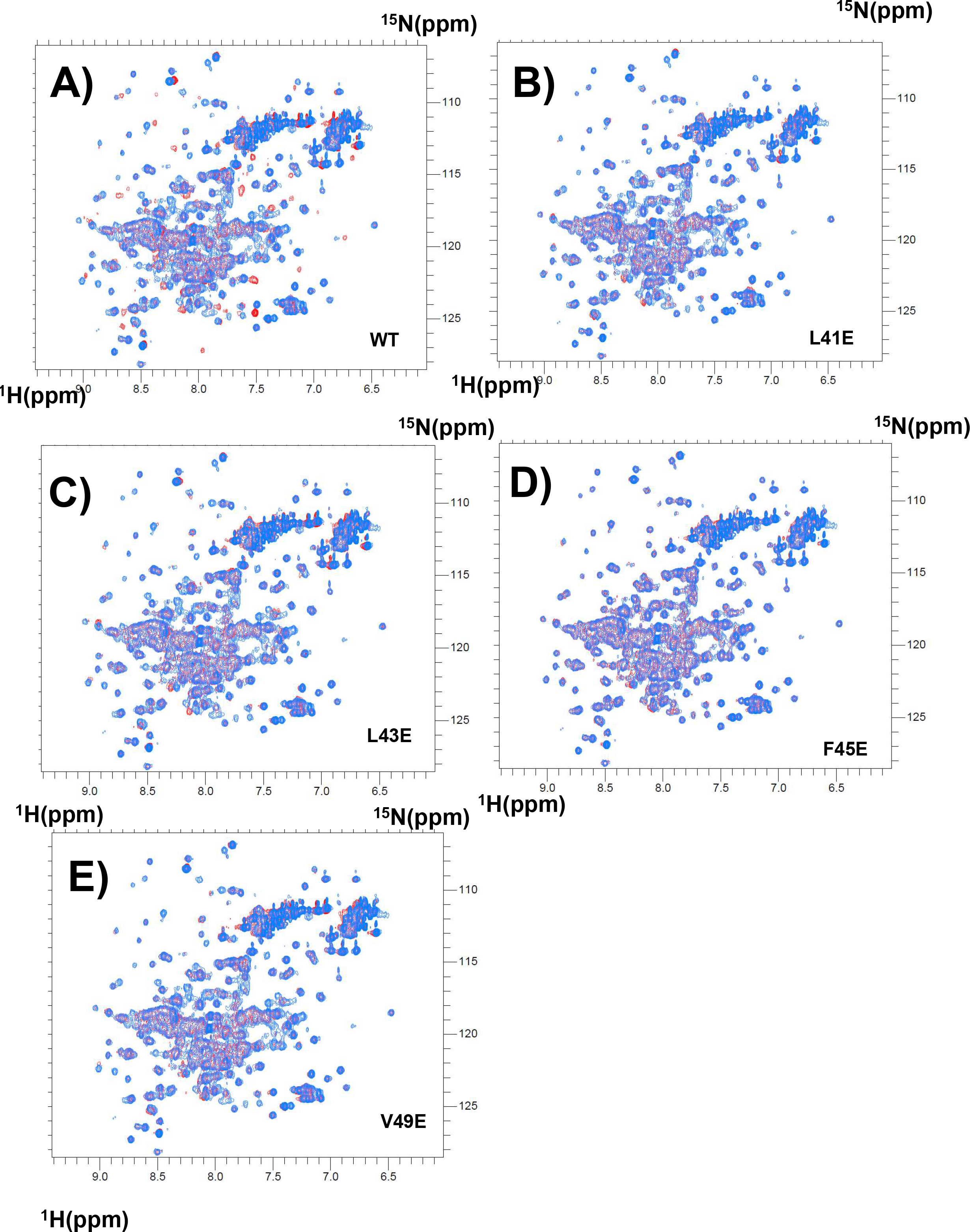
Full spectra of 400 μM ^1^H,^15^N-labelled R7R8 with synthetic KANK1 peptides containing mutations **A)** WT, **B)** L41E, **C)** L43E, **D)** F45E **and E)** V49E. Blue peaks represent control spectra and the addition of KANK1 peptide is red.

**Figure S7.**
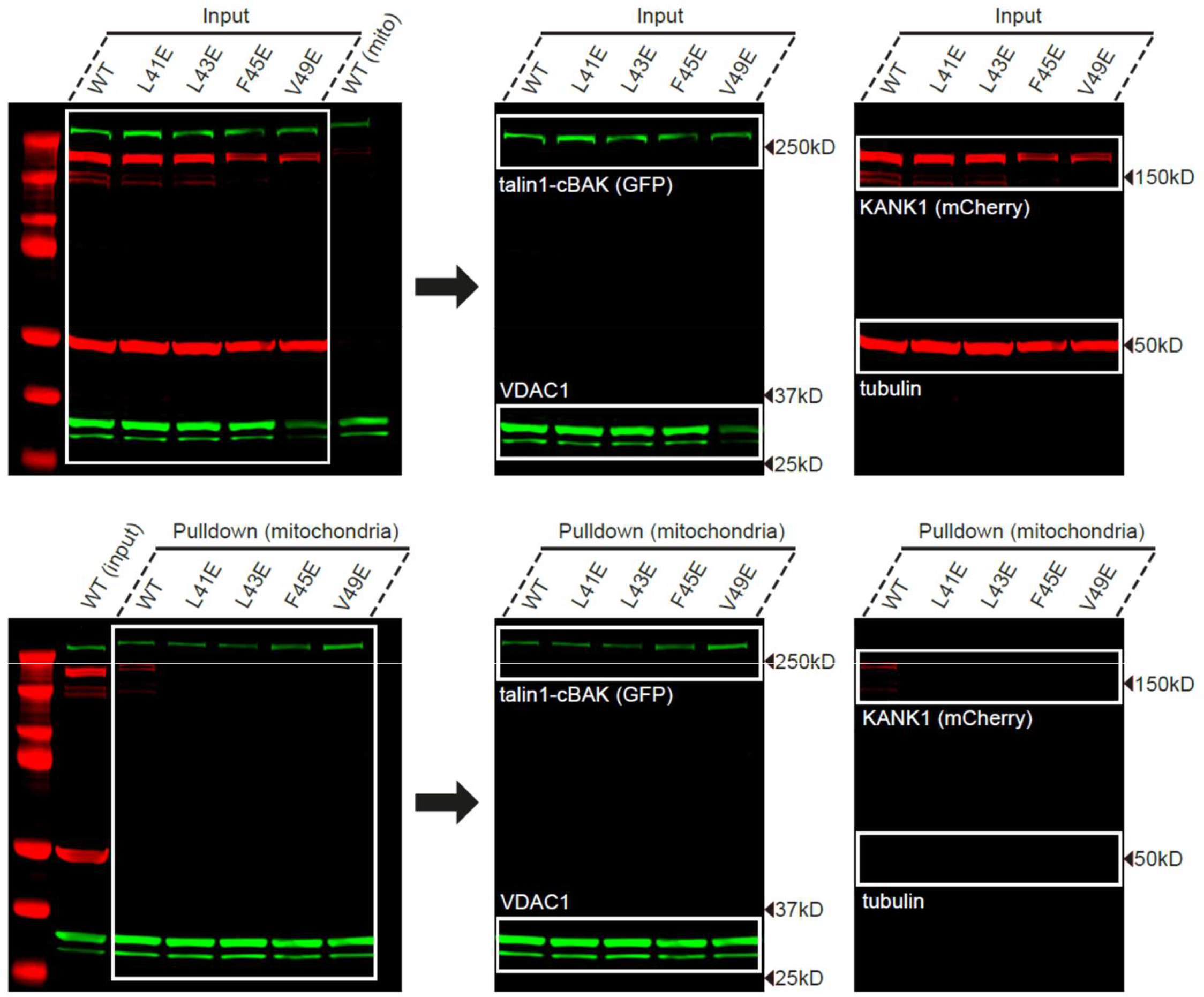
Raw western blots and repeats of cBAK colocalization assay experiments with input proteins on the top row and immunoprecipitation experiments below.

**Figure S8.**
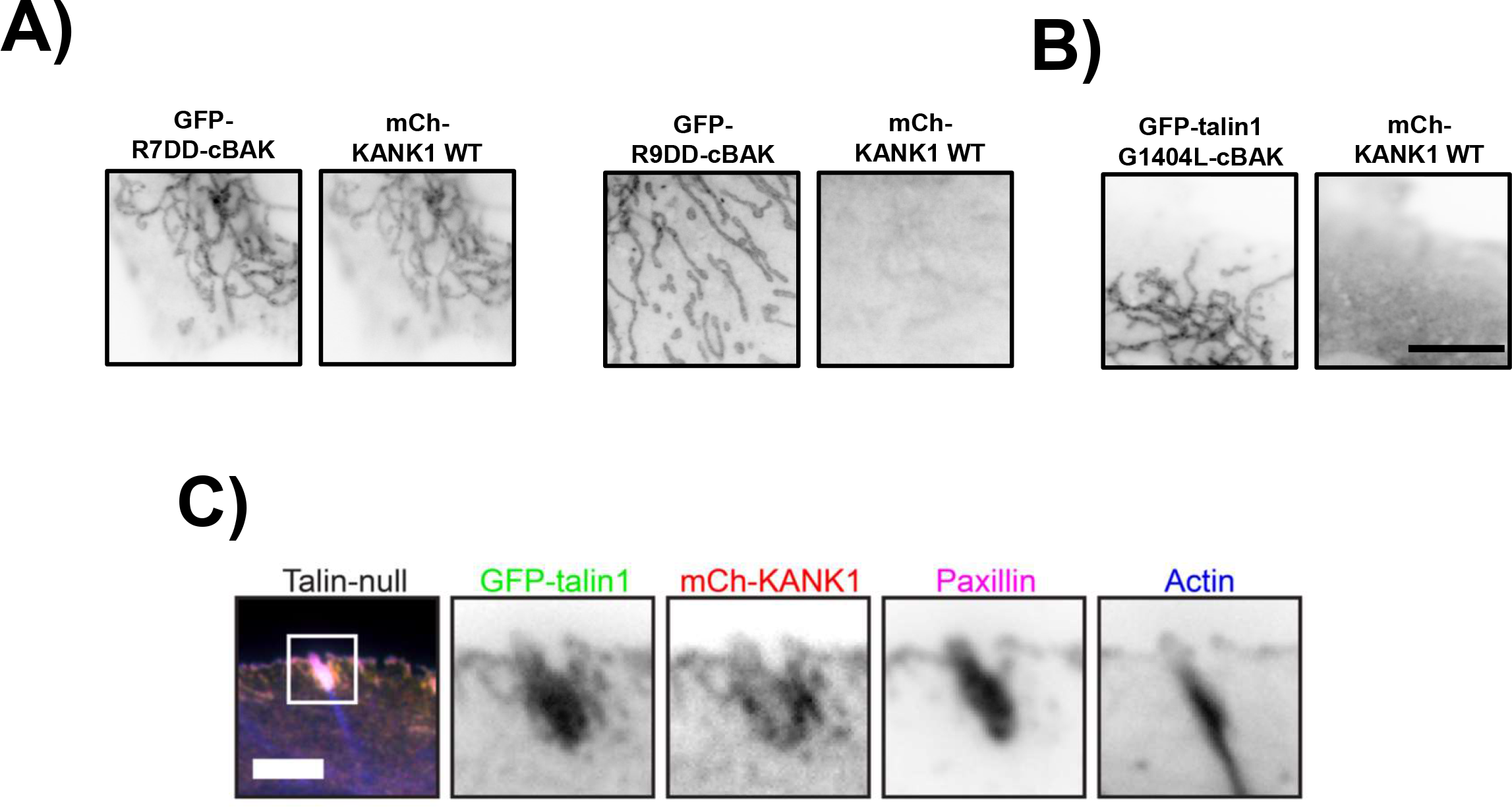
Mitochondrial targeting assays of **A)** GFP-R7DD-Cbak with mCherry-KANK1, **ii)** GFP-R9DD-Cbak with mCherry-KANK1. **B)** GFP-talin1 G1404L-Cbak with mCherry-KANK1. **C)** Line distribution profile of adhesions containing GFP-talin-1, mCherry-KANK1 stained for actin and paxillin in TKO cells. Scale bar is 5μm.

**Figure S9.**
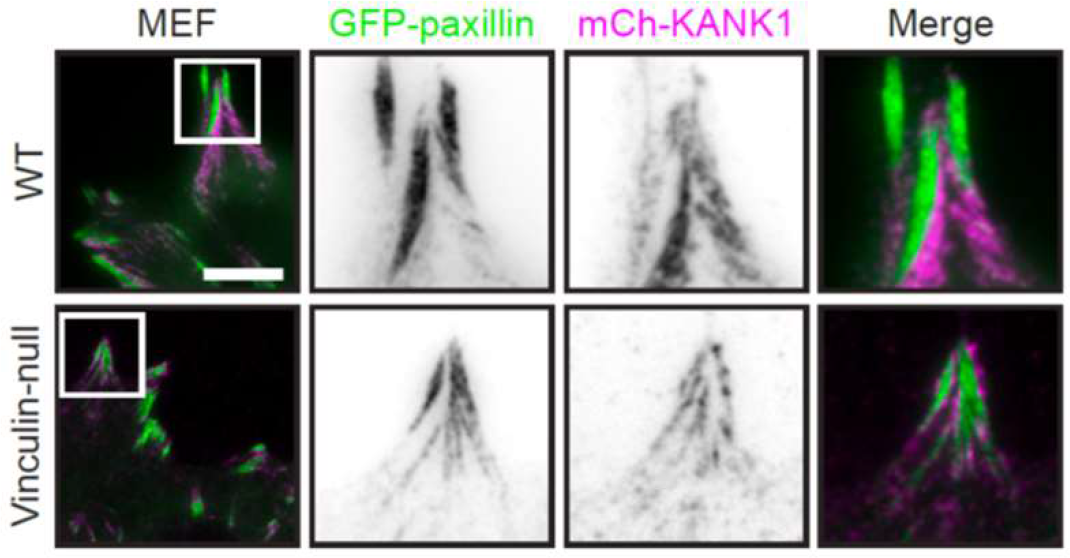
Images of adhesions in WT and vinculin-null MEFs expressing GFP-paxillin and mCherry-KANK1. Scale bar 10 μm.

**Figure S10.**
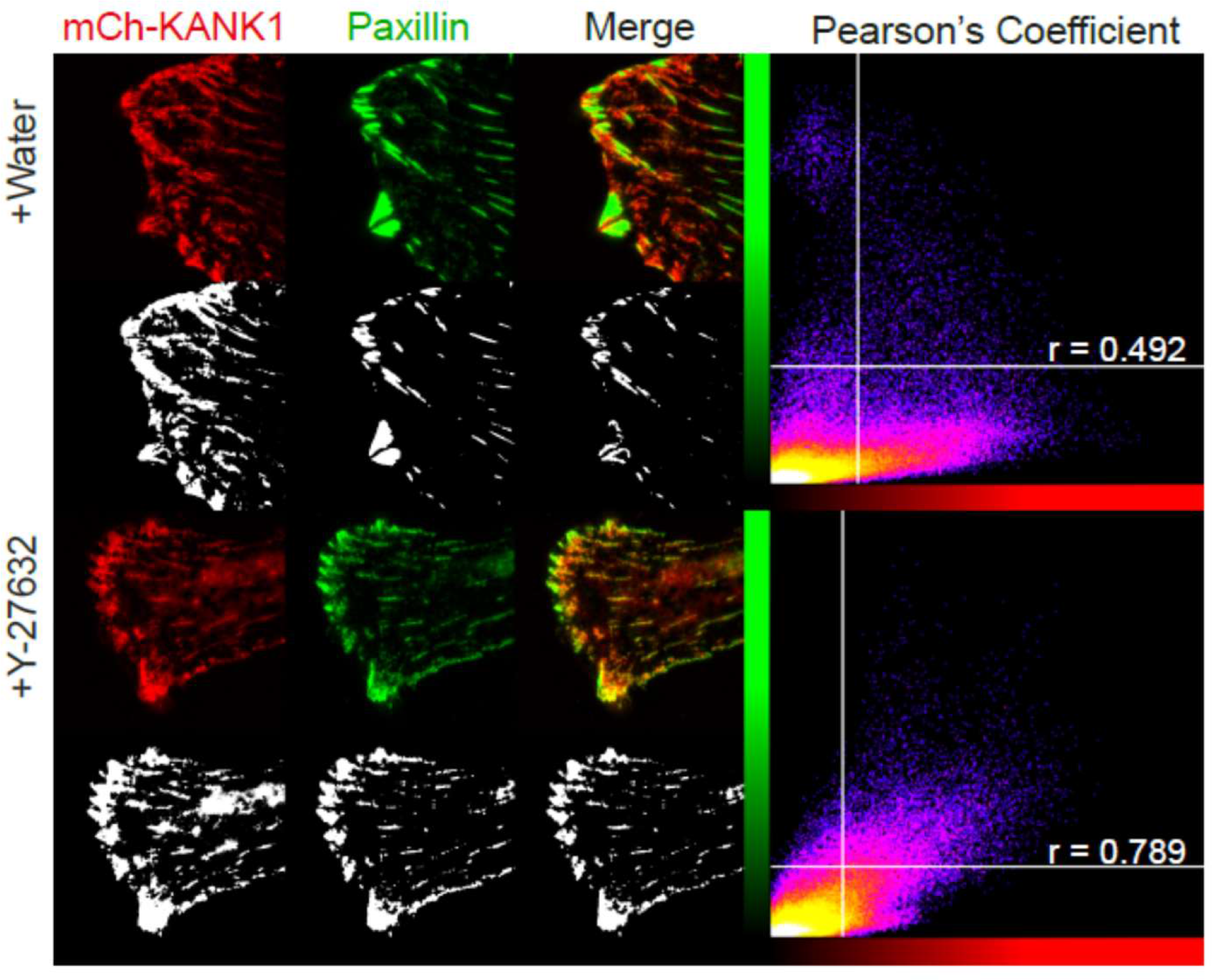
Pearson’s correlation coefficient quantification of 20 (n) individual 40*40 μM regions, where KANK1 and paxillin overlap in control cells with R=0.59, and in treated cells R=0.82.

